# *In utero* pulse injection of isotopic amino acids quantifies protein turnover rates during murine fetal development

**DOI:** 10.1101/2023.05.18.541242

**Authors:** Josue Baeza, Barbara E. Coons, Zongtao Lin, John Riley, Mariel Mendoza, William H. Peranteau, Benjamin A Garcia

## Abstract

Protein translational control is highly regulated step in the gene expression program during mammalian development that is critical for ensuring that the fetus develops correctly and that all of the necessary organs and tissues are formed and functional. Defects in protein expression during fetal development can lead to severe developmental abnormalities or premature death. Currently, quantitative techniques to monitor protein synthesis rates in a developing fetus (*in utero*) are limited. Here, we developed a novel *in utero* stable isotope labeling approach to quantify tissue-specific protein dynamics of the nascent proteome during mouse fetal development. Fetuses of pregnant C57BL/6J mice were injected with isotopically labeled lysine (Lys8) and arginine (Arg10) via the vitelline vein at various gestational days. After treatment, fetal organs/tissues including brain, liver, lung, and heart were harvested for sample preparation and proteomic analysis. We show that the mean incorporation rate for injected amino acids into all organs was 17.50 ± 0.6%. By analyzing the nascent proteome, unique signatures of each tissue were identified by hierarchical clustering. In addition, the quantified proteome-wide turnover rates (k_obs_) were calculated between 3.81E-5 and 0.424 hour^-1^. We observed similar protein turnover profiles for analyzed organs (*e.g.*, liver versus brain), however, their distributions of turnover rates vary significantly. The translational kinetic profiles of developing organs displayed differentially expressed protein pathways and synthesis rates which correlated with known physiological changes during mouse development.

## INTRODUCTION

During development before birth, multicellular, complex organisms differentiate into many different cell types from a single zygote in a highly ordered and reproducible manner. Precise spatial and temporal regulation of the gene expression program is critical for normal development^1^. Protein biosynthesis, the translation of mRNA into protein, is an essential step in gene expression and has the highest energy demand in the cell^2, 3^.

The translational machinery, originally considered a “housekeeping” function, has emerged as a highly regulated and dynamic process in gene expression. Translational regulation allows for more rapid changes in the proteome, allowing cells to respond to external stimuli^4, 5^, differentiate^6, 7^, and maintain homeostasis^6–8^. Recently, translational regulation, for global and specific mRNA transcripts, has been described as a regulatory mechanism in itself^9^. For example, stem cells maintain a low basal translational output including hematopoietic stem cells^10^, neural stem cells^11^, muscle stem cells^12^, epidermal stem cells^13^, and *Drosophila* germline stem cells^14^. Consequently, dysregulation of translational control, through pharmacological or genetic means, leads to impaired stem cell function. Therefore, understanding the mechanisms of translational control regulating cell function and animal development requires quantification of protein turnover *in vivo*.

Proteins are in a dynamic state of equilibrium, continuously being synthesized and degraded^15^. Protein turnover is the combined action of protein synthesis and degradation and is responsible for establishing and modulating protein abundance levels in cells^5^. Protein half-lives are linked to protein function and display a wide range: from minutes (∼20 min for p53 in humans)^16^ to an organism’s entire lifetime (>70 years for the eye lens crystallin in humans)^17^. Despite its importance in cell function and differentiation^6^, protein turnover has not been systematically investigated in the context of fetal development.

Quantification of *in vivo* mammalian protein turnover requires the introduction of stable isotope-labeled amino acids, usually through the diet^18, 19^. The time needed to quantify protein turnover rates requires weeks of isotope labeling, which is outside the scope needed to capture changes in protein synthesis during murine development as well as in other animals with short gestation periods. Additionally, isotopic labeling via the diet is somewhat expensive due to having to label the animal for long periods (e.g. weeks). In this study, we developed a novel pulse injection method to label proteins in the developing fetus at defined gestational periods. Unique protein turnover rates were determined at various days of gestation, providing a high temporal resolution of turnover rates in the developing mouse. Our approach utilizes a direct injection of stable isotope-labeled amino acids into the fetal circulation which are taken up by tissues and utilized for protein synthesis. NanoLC-MS/MS analysis of the liver, heart, and lungs from pulse-injected fetuses quantifies the nascent and pre-existing proteome of the developing organs. The small size of the fetus allows the injectate to be maximized for proteome incorporation. By sampling organs at defined time intervals after the injection, we showed that protein turnover rates could be quantified and that changes in the synthesized proteins correlate with known physiologic changes in the fetal liver during mouse development.

## RESULTS

### Pulse injection for *in utero* labeling

We sought to evaluate the nascent proteome in the fetus and changes to the proteome, including protein turnover rates, in the context of normal ontologic changes during fetal development. As such, we developed a method, in the murine model, of direct fetal intravascular injection of labeled amino acids to exclude the requirement of maternal-fetal transfer of maternally fed or injected labeled amino acids and to precisely time the exposure of the fetus to the labeled amino acids with subsequent downstream analyses. To label the nascent proteome *in utero*, fetal mice are pulse-injected via the vitelline vein with a bolus of isotopic amino acids^20, 21^. The vitelline vein, which runs along the uterine wall, drains directly into the hepatic portal circulation resulting in a first-pass effect on the fetal liver. The injected labeled amino acids are taken up by the developing organs and used by the translational machinery for protein synthesis. The incorporation of heavy-isotope labeled amino acids into the proteome can then be quantified by high-resolution mass spectrometry (**Fig. 1**).

**Figure 1:**
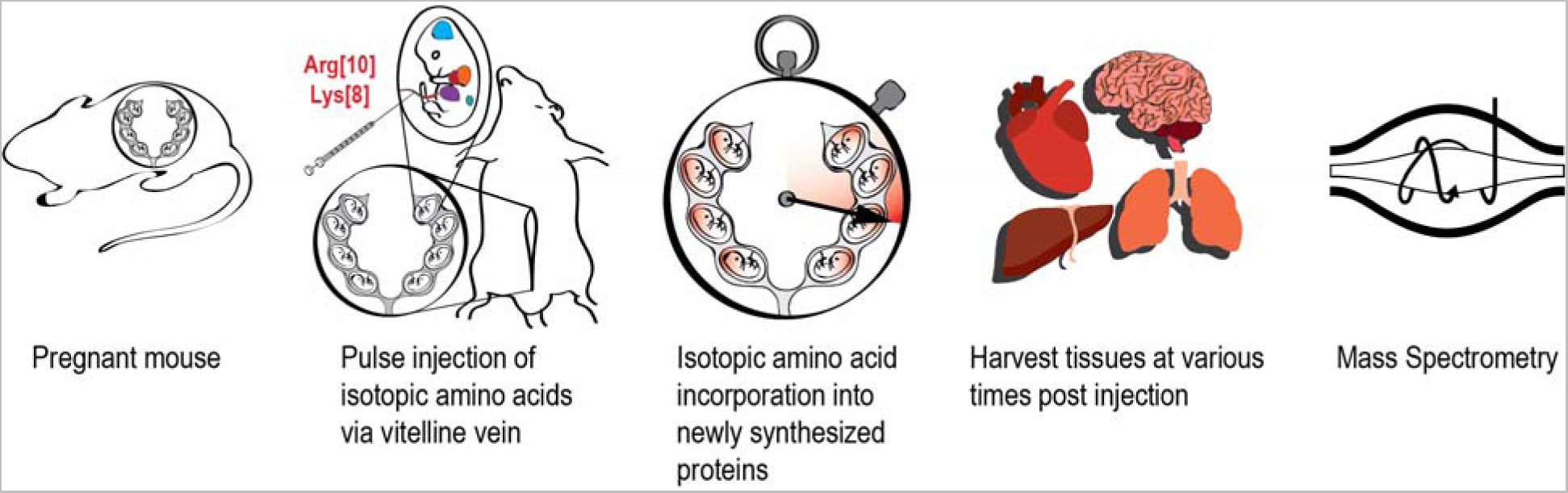
Pulse-injection method for *in utero* protein labeling. A time-dated pregnant mouse undergoes laparotomy under isoflurane anesthesia to expose the uterine horns. 10 μmol of stable isotope-labeled L-lysine and L-arginine in sterile PBS was injected into each fetus through the uterine wall via the vitelline vein. After injection, the fetuses were returned to the peritoneal cavity and the surgical incision was sutured. Pregnant dams recover after 15-20 min under a heat lamp. Labeled amino acids circulate throughout the fetus and are transported to developing organs to be used for protein synthesis. At specific time points after amino acid injection, fetal tissue is harvested and analyzed by high-resolution mass spectrometry.

To determine the extent of incorporation of the exogenous amino acids into the proteome, gestational day (E) 14.5 fetal mice (normal gestation is 20 days) were pulse-injected with isotopic amino acids and organs were harvested (liver, heart, and lung) 16 hours post-injection and analyzed by high-resolution mass spectrometry-based proteomics (**Fig. 2A-B** and **Supplementary Dataset 1**). The mean incorporation rate (heavy fractional abundance) into all organs analyzed was 17.50 ± 0.6%, (median incorporation 8.6 ± 1.2%, **Fig. 2C**). We observed very high reproducibility across different organs and pulse injection replicates (**Fig. 2D**). Light (non-isotopic, endogenous) and heavy labeled peptides showed the highest correlations when comparing pulse-injection replicates of the same organ (Pearson correlation coefficient > 0.95). Measurement variability of light and heavy peptide abundances was low with the median coefficient of variation (CV) values at 13.80% and 21.97%, respectively (**Fig. S1**). Principal component analysis (PCA) performed on the peptide features identified distinct groups for the light and heavy proteomes which were also grouped by tissue. The first principal component represents the pre-existing (light) and nascent (heavy) proteomes, while the second principal component represents different tissues (**Fig. 2E**). The nascent proteome was approximately an order of magnitude lower than the pre-existing proteome (**Fig. 2B**). We compared the abundance distribution of the nascent proteome and found it to be a subset of the most abundant peptides present in the total proteome sample (**Fig. 2F**). From these data, we determined that a 16 hour labeling period is sufficient to label the distinct tissue proteomes and a subset of proteins being actively synthesized in different organs of a developing mouse fetus.

**Figure 2:**
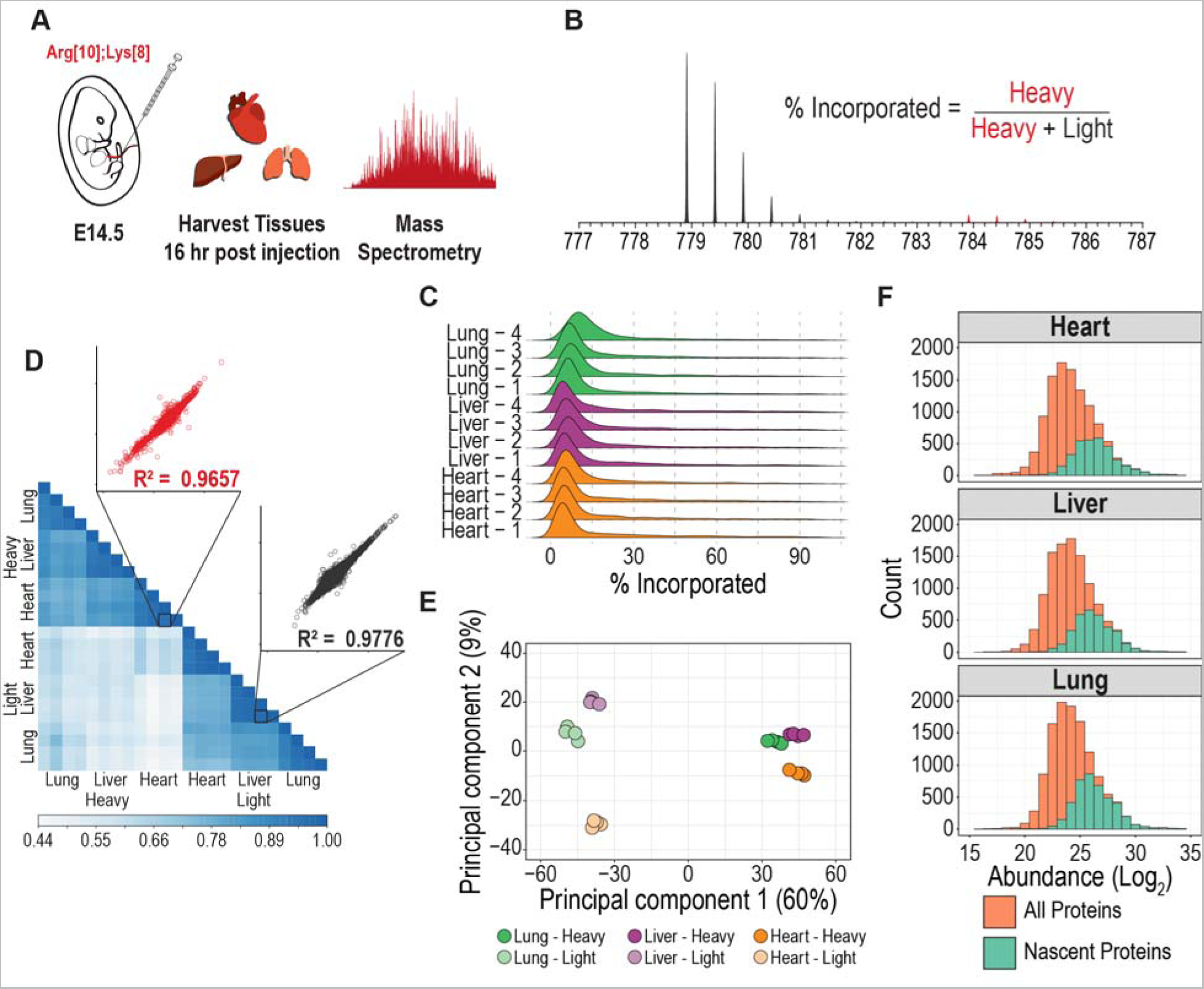
Method development of pulse injection for fetal labeling. **A.** Experimental design for *in utero* labeling. Fetuses from E14.5 pregnant mice are pulse-injected. Heart, liver, and lung tissues are harvested 16 hours later and analyzed by mass spectrometry. **B.** Representative MS1 spectra showing heavy amino acid incorporation. % incorporation is calculated as the fraction of heavy precursor over the total precursor pool (heavy + light). **C.** Incorporation of heavy labeled amino acids into the proteome of heart, liver, and lung tissues using four biological replicates. The median and mean incorporation is 8.6 ± 1.2% and 17.5 ± 0.6%, respectively. **D.** Heatmap showing the Pearson correlation of the light and heavy precursors across the tissues analyzed. Inset: Scatter plots of light and heavy peptides show high reproducibility between biological samples. **E.** Principal component analysis (PCA) of the light and heavy precursors. **F.** Distribution of peptide abundances in Log_2_ showing the subset of nascent peptides (heavy-isotope labeled).

We next examined whether pulse injection of heavy amino acids followed by different labeling periods would result in the quantification of the nascent proteome. For this, E14.5 fetuses were pulse-injected with labeled amino acids and the heart and liver were harvested for proteomic analysis at 8, 24, and 48 hours post-injection (**Fig. S2A**). Hierarchical clustering of the nascent proteome identified a unique signature for each tissue (**Fig. S2B**). Gene Ontology (GO) analysis of the enriched clusters show enrichment of terms associated with “development” for the heart, while the liver shows GO term enrichment associated with “translation”. These results are in line with the function of these organs at this stage of development. At gestational age E14.5 to E16, the liver is the major site of hematopoiesis and, among other functions, supports hematopoietic stem cell development and differentiation including cells responsible for hemoglobin synthesis^22^, while the heart completes the development of the atrial septation.^23^

The incorporation of heavy amino acids into the proteome at 8, 24, and 48 hours showed similar distribution profiles as the 16-hour labeling period (**Fig 2C and Fig. S2C**). However, the similar incorporation profiles may not reflect differences in individual proteins’ heavy abundances across different labeling periods. To test this hypothesis, we performed a statistical analysis on the nascent proteome to compare the abundances across the different labeling periods (**Fig. S2D**). An enrichment of GO terms associated with “binding of sperm to zona pellucida” and “regulation of establishment of protein localization to telomere” was observed in the liver while the heart showed enrichment in GO terms associated with “sperm-egg recognition” (**Fig. S2E**), GO terms associated with development. Taken together, we demonstrate that the proteome of fetal organs can be labeled *in utero* with isotopic amino acids to distinguish between the nascent and pre-existing proteome and this approach can quantify changes in the nascent proteome using varying labeling periods. However, using these labeling periods, the quantification of the nascent proteome (heavy) represents the cumulative effects of multiple cellular processes such as translation, degradation, secretion, and even cell division. Disruption of any of these processes (*e.g.,* proteasome inhibition) would affect the observed labeling efficiency. To quantify protein turnover rates, shorter labeling periods would be required.

### Quantifying protein turnover rates *in utero*

To understand how changes in protein turnover regulate normal tissue development and homeostasis, it is important to quantify proteome-wide protein turnover rates *in utero*^9^. To quantify turnover rates, we utilized short labeling periods (1, 2, 4, and 6 hours) (**Fig. 3A**). In this experimental design, injection of the labeled amino acids corresponds to a quick pulse followed by the chase with endogenous amino acids, which are sourced by the maternal diet. The distribution of heavy (isotopically labeled) fractional abundance (H / (H + L)) increases with the labeling period (**Fig. S3A-B**). The principal component analysis also revealed a natural progression of the heavy-isotope labeled proteome during the time course for each tissue (**Fig. 3B**). Turnover rates were determined by fitting the heavy fractional abundance of each peptide to an exponential decay model^24–27^ which quantified turnover rates (k_obs_). As an example, we show the turnover rate of the 40S ribosomal protein S12 (RPS12) to be 0.0804 hour^-1^ and 0.227 hour^-1^ in the liver and brain, respectively (**Fig. 3C, 3D**). The quantified proteome-wide turnover rates (k_obs_) ranged between 3.81E-5 and 0.424 hour^-1^, which equates to half-lives ranging between 1.6 hours to >500 days. Proteins displaying extremely long half-lives include histones, which are known to be long-lived proteins^28^.

**Figure 3:**
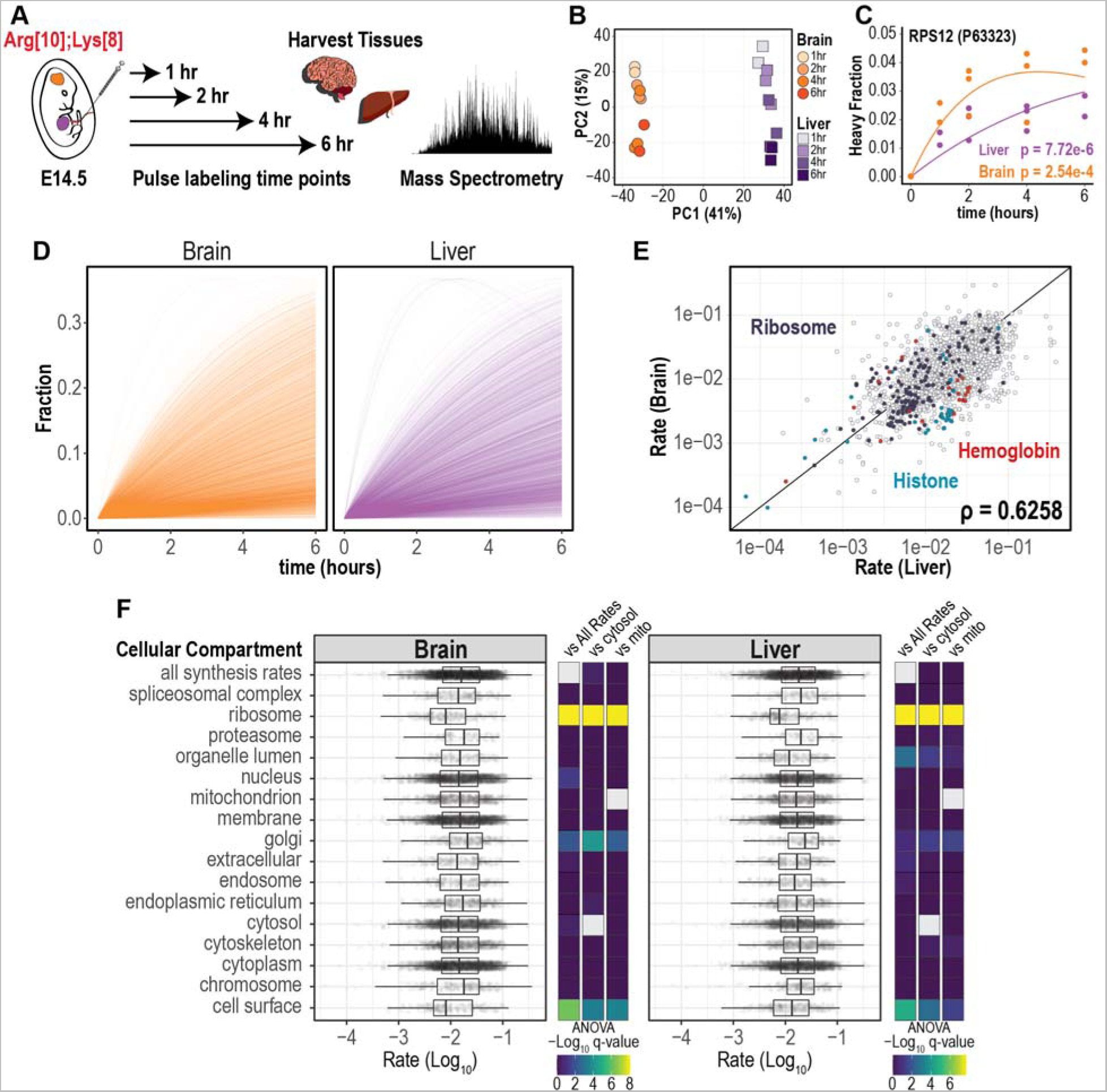
Quantifying *in utero* protein turnover by pulse-injection. **A.** Experimental design for quantifying protein turnover. Fetuses from E14.5 pregnant mice are pulse-injected with labeled amino acids. Fetal liver and brain are harvested from different pregnant mice at 1, 2, 4, and 6 hours after injection and analyzed by mass spectrometry. **B.** Principal component analysis for the heavy peptides from brain and liver tissues at 4 time points are shown. **C.** Modeling the heavy peptide fraction as a function of time using a non-linear pulse-chase turnover model. The data is from the 40S ribosomal protein S12 (RPS12) and the p.values of the model fits are shown. **D.** Turnover rate profile across the proteome of brain and liver tissues. **E.** Spearman correlation of turnover rates between brain and liver tissues. Peptides corresponding to Ribosome, Histone, and Hemoglobin proteins are colored. **F.** Distribution of log_10_-transformed turnover rates grouped by cellular compartment. Cellular localization was performed in Proteome Discoverer 2.2 and each data point corresponds to a single peptide turnover rate. Analysis of variance (ANOVA) followed by Benjamini-Hochberg correction was used to compare turnover rate distribution between cellular compartments and all rates (vs All Rates), cytosol (vs Cytosol), and mitochondria (vs Mito).

We also quantified the labeling kinetics of the precursor amino acid pool. For this, we measured the fractional abundance of the heavy amino acid pool and fit the data to an exponential decay model as a function of labeling time. To monitor the amino acid precursors, we used peptides with two lysines or two arginines and performed mass isotopomer distribution analysis (MIDA) to determine the distribution of the heavy amino acid pool^18, 29–31^ (**Fig. S3C**). The median fractional abundance of each sample was used to fit the exponential decay model (**Fig. S3D**). Based on this analysis, the half-life of the amino acid pool in the brain and liver is 6.1 and 5.5 hours, respectively, which is within the labeling period of the experiment.

Next, we compared protein turnover rates for the liver and brain and observed similar profiles for these tissues (**Fig. 3D**). In this analysis, each line represents a single model fit, which represents all of the time points and replicates. We then compared the distribution of turnover rates between the liver and brain and found a significant difference between the two profiles. This difference is mediated by a different distribution of proteins with low turnover rates (**Fig. S4A**). Synthesis rates between the brain and liver displayed a moderate correlation (*p* = 0.6258) (**Fig. 3E** and **Supplementary Dataset 2A-B**), however, distinct protein families showed higher rates in different tissues. For example, hemoglobin and histone proteins had a higher turnover rate in the liver, highlighting the differences in tissue function at the E14.5 stage of development. Specifically, the fetal liver is the predominant site of hematopoiesis at E14.5 at which time hematopoietic cells are undergoing rapid expansion^32, 33^. To identify proteins with significant changes in turnover rate between the liver and brain, we performed a Mann-Whitney test followed by correction for multiple hypothesis testing. Proteins with a higher turnover rate in the liver included Alpha-Fetoprotein, Lamin-B1, Pyruvate Kinase PKM, and Hemoglobin β1. Proteins with a higher turnover rate in the brain included leucine-tRNA synthetase, ribosome binding protein 1, adenosylhomocysteinase, and E3 ubiquitin ligase NEDD4 (**Fig. S4B**). We also identified protein abundance to be negatively correlated with the turnover rate (**Fig. S4C**). This observation has been reported previously^18^ and suggests that the cell is balancing energetic costs and protein turnover.

Differences in subcellular turnover rates were also observed. For this analysis, we categorized proteins based on known subcellular location and compared turnover rates between the various subcellular regions (**Fig. 3F**). Ribosomal proteins had significantly lower turnover rates in the liver and brain. Proteins residing in the Golgi and cell surface also displayed significant differences when compared to other cellular regions. Proteins that are annotated as “nucleus”, “membrane” and “cytoplasm” displayed a uniform distribution. Taken together, this method can be used to quantify *in utero* turnover rates using labeling periods as little as 1 hour to 6 hours. This labeling period is within the period of cell division, therefore cell growth is not a major confounder of the turnover rate measurement.

### Quantifying turnover rates across fetal development

We next sought to understand how protein turnover rates change across fetal development. To do this, we pulse-injected fetal or neonatal mice via the vitelline vein or facial vein^34^, respectively, with heavy labeled amino acids (Lys8 and Arg10) at E13.5, E14.5, E16.5 and day of life 0 (P0). Livers and lungs from injected mice were harvested at 2, 4, and 6 hours post-injection to provide respective labeling time points for analysis (**Fig. 4A, Fig. S5A**). To compare between the different organs, labeling time points and developmental stages at the time of injection, we included a pooled liver reference sample as a standard to run with each sample batch. We also included a pooled lung reference sample, which was only analyzed with the lung samples. The pooled liver-std reference sample was used to harmonize the datasets between liver and lung samples using a single-point calibration method (**Fig. 4B**)^35^. To validate the data harmonization, we used a Uniform Manifold Approximation and Projection (UMAP) for dimensionality reduction and visualization. The samples show a clear separation between the two tissues as well as by developmental stages (**Fig. 4C** and **Supplemental Dataset 3**). Additionally, the pooled reference samples cluster with their respective tissue. Another layer of validation is to identify proteins with known biological functions at defined gestational periods in development. The liver is the predominant site of hematopoiesis at E14.5-producing cells that constitute the lymphohematopoietic system including the erythroid lineage^32^. Therefore, we quantified the turnover rates of hemoglobin proteins in the liver and noted that hemoglobin α and β subunits displayed peak turnover rates at E14.5 with the rate decreasing at later gestational ages (**Fig. 4D, Fig. S5C, Supplemental Dataset 4**).

**Figure 4:**
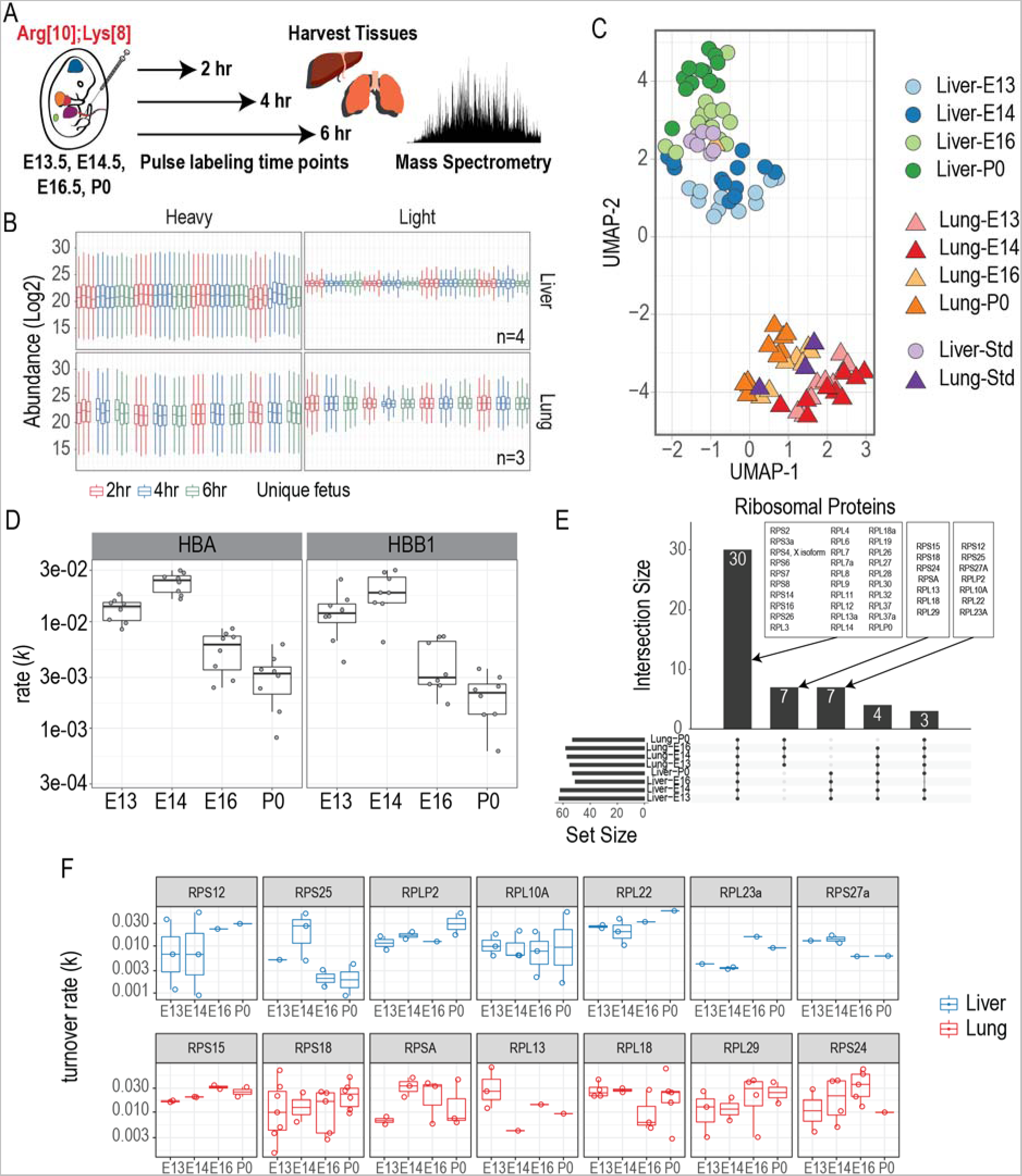
Quantifying *in utero* protein turnover across mouse fetal development. **A.** Experimental design for quantifying protein turnover. Fetuses from E13.5, E14.5, and E16.5 are pulse-injected with labeled amino acids. P0 mice are pulse-injected through the retro-orbital vein^34^. Fetal liver and lung tissues are harvested from different pregnant mice at 2, 4, and 6 hours post-injection and analyzed by mass spectrometry. **B.** Distribution of batch-corrected and normalized log_2_ transformed peptide abundances. MS runs were batch-corrected using a common standard protein sample that was processed with every batch. **C.** UMAP dimensionality reduction of the liver and lung proteomes from all time points. **D.** Turnover rates of hemoglobin proteins in the liver at different gestational ages. **E-F.** Quantified peptide turnover rates for different proteins at defined gestational stages from the liver and lung, respectively.

Gene expression requires many regulatory layers to convert the genetic information found in DNA to phenotype. Emerging evidence has linked ribosome heterogeneity to gene regulation^36^. We sought to determine if ribosome heterogeneity and translational control might be observed with protein turnover of developing liver and lung tissues. Ribosomal proteins with quantified turnover rates were grouped based on tissue profiling (**Fig. 4E**). We quantified translation rates for 30 ribosomal proteins during development which were common to both liver and lung tissues. Additionally, we quantified turnover rates for ribosomal proteins that were unique to each tissue with varying trends (**Fig. 4E, 4F**). 40S ribosomal protein S25 (RPS25) and 60S ribosomal protein L10A (RPL10A) were quantified in the liver during all developmental stages analyzed. RPS25 displayed peak levels in the liver at E14.5, while RPL10A displayed a constant rate during development. Interestingly, RPS25 and RPL10 are involved in IRES-dependent translation ^37,38^suggesting that these proteins might be involved in translating distinct mRNA pools. We also quantified turnover rates for the 40S ribosomal protein S24 (RPS24) in the lung. The turnover rate of RPS24 increased from E13.5 to E16.5 with a sudden decrease at P0. Mutations in RPS24 have been linked to Diamond-Blackfan anemia^39^ (see discussion). Further studies will be needed to understand how specialized ribosomes and their turnover rates are involved in fetal development.

## DISCUSSION

In this study, we developed an approach utilizing *in utero* pulse injection of isotope-labeled amino acids to quantify protein turnover rates in a developing mouse fetus. Turnover rates for individual proteins represented a wide range across tissues and developmental stages. Changes in turnover rates also coincided with known physiologic functions of tissues at defined developmental stages. There are many benefits of using the *in utero* pulse-injection method for determining turnover rates include: First, stable isotope labeled amino acids are introduced directly into the fetal circulation, avoiding the long labeling periods required for providing labeled amino acids via the diet^18^. The time required to label a mouse via the diet is usually at least a few days^24, 40^, which is a significant portion of time during the normal gestational development of a mouse, highlighting the inability of using this method to study fetal protein turnover rates during gestation. Secondly, the mode of injection via the vitelline vein results in the first-pass effect of the labeled amino acids in the fetal liver with the subsequent systemic distribution of the amino acids to the rest of the fetus. This allows the dose injected into each fetus to be efficiently incorporated into the proteome. Thirdly, the amount of isotopic amino acids is minimized, therefore reducing the costs associated with each experiment. The amount of isotopic amino acids used for a 0.1 g fetal mouse^41^ was 10 μmol compared to the required 3.5 grams of heavy-isotope labeled chow needed for a 20 g adult female mouse^19^. Finally, the labeling period can be performed with minimal incubation time (1-6 hours) to provide a high temporal resolution of protein turnover during mouse development. In the present study, we quantified turnover rates at four gestational time points of mouse fetal development.

Amino acid analogs containing an azide moiety, such as azidohomoalanine (AHA), can be incorporated into nascent proteins during translation and subsequently analyzed by mass spectrometry^42^. Using our treatment method, we did not observe significant proteome differences between AHA and control (no injection) in three major organs (**Fig. S6**). This is reasonable since natural amino acids, whenever available, will be preferably utilised for protein synthesis compared to their non-natural counterparts (*e.g.*, AHA). AHA treatment is a attractive enrichment strategy to profile the nascent proteome and allow the quantification of low-abundant proteins. This strategy, combined with stable isotope labeling, has enabled the study of the nascent proteome^43, 44^. However careful consideration must be taken when using AHA to profile the embryonic proteome because, if incorporated, the non-natural amino acid at this early stage of development has the potential to impact the proteome^45, 46^.

Comparing the proteomes of three organs from mice received saline treatment and control (no injection), we show that the injections themselves did not cause major expression changes during the fetal development (**Fig. S6**). The dose administered was the same (10 μmol) for all fetal injections, regardless of embryological age. As a result, there may be some variation in the ratio of isotopic amino acids to fetal weight. However, we observed no significant difference in the incorporation rate of the isotopic amino acids as a function of gestational age. If desired, the *in utero* pulse injection technique could be adapted to administer a dose that is proportional to the size of the fetus for each gestational age. With new tools to quantify turnover dynamics of the nascent proteome *in utero*, fundamental questions about how changes in protein turnover regulate cell function, and fetal development or cause clinical disorders can be addressed^9^, which would have a broad effect on embryology and developmental biology. For example, this method could be used to study the effects on the fetal translatome from a diverse array of exposures. such as mouse models of intrauterine growth restriction, chorioamnionitis, and twin-twin transfusion syndromes. Changes in fetal protein turnover or relative protein abundances could then be used to understand phenotypic outcomes.

Ribosomopathies are characterized as disorders caused by the reduced expression of, or mutations in ribosomal proteins or ribosome biosynthesis factors, which manifest in human disease with diverse tissue-specificity^47^. Because of the translational machinery’s critical role during development, ribosomopathies are classified as developmental disorders. Diamond-Blackfan Anemia (DBA) manifests with bone marrow failure and is caused by mutations in the 40S ribosomal protein S19 (RPS19) ^48^, RPS24^39^ as well as other ribosomal proteins. The mechanism of DBA pathogenesis remains unresolved^49^. Our method of quantifying *in utero* protein turnover rates could be applied to potentially identify targets of RPS19 or RPS24 causing DBA. Understanding the mechanisms of DBA pathogenesis can lead to better treatment options for patients.

## LIMITATIONS OF THE STUDY

One of the limitations of this approach is the limited dynamic range of proteins profiled due to the narrow incorporation rate of isotopic amino acids into the proteome (∼10%). For this reason, highly abundant proteins were the majority of proteins with quantified turnover rates. However, various strategies can be employed to overcome this limitation. For example, multiplexing various time points and including a “booster” channel will allow high sensitivity and accurate quantification^5^ especially when combined with offline fractionation. Alternatively, high-sensitivity methods such as data-independent acquisition (DIA) can be utilized to extend the dynamic range of the proteins profiled as we have recently shown to quantify cellular protein turnover^50^.

During embryonic development, non-dividing cells of a specific organ/tissue may utilize the injected amino acids at lower rate than dividing cells which require free amino acid for protein synthesis. The protein turnover reported in this study focused on the overall tissue/organ level, thus does not necessarily represent the protein synthesis of dividing cells from each tissue/organ.

It is reported that Arg10 can be metabolically converted to [^13^C_5_,^15^N_1_]-proline resulting in a Δmass of 6 Da with unlabeled proline^51, 52^. Peptides containing such resulted proline may subject to metabolic bias in protein turnover. Since metabolic conversion happens systematically across all the tissues, for simplicity of the method development, we did not take such conversion into consideration during the data analysis.

## CONCLUSION

In this study, a novel *in utero* approach has been established to monitor the amino acid usage during various mouse fetal developmental stages. The method enables quantification of the nascent proteome in a tissue-specific manner thus allows for the measurements of translational protein kinetics. To the best of our knowledge, this is the first time that an *in utero* method has been developed to evaluate the protein turnover during fetal development. The pulse-injection strategy of isotopic amino acid has demonstrated success in differentiating the existing versus the newly synthesized proteomes in different organs/tissues of a developing mouse fetus. In our efforts to quantifying protein turnover rates *in utero*, we have obtained data to understand how these rates change across feta development stages (*e.g.*, E13.5, E14.5, E16.5 and P0). Having precise temporal control of fetal labeling will allow one to investigate translational control and regulation throughout fetal development. The ability to study the fetal proteome at such a level will lend insight into developmental biology and the evolution of pathologic processes.

## STAR METHODS

Detailed methods are provided in the online version of this paper and include the following:

- KEY RESOURCES TABLE
- RESOURCE AVAILABILITY

`o` Lead contact
`o` Materials availability
`o` Data and code availability
- METHOD DETAILS

`o` Animals
`o` *In utero* and neonatal pulse-injection of labeled amino acids
`o` Sample processing
`o` LCMS analysis
`o` Data analysis
`o` Mass isotopomer distribution analysis
`o` Determination of turnover rate
`o` Statistical analysis

## SUPPLEMENTARY INFORMATION

1. Proteomics data of three fetal organs (liver, heart, and lung) after pulse-injection of isotopic amino acids at E14.5 stage. (**1 csv file**)
2. Proteomics data on protein turnover in two organs (liver and brain). (**7 csv files**)
3. Proteomic data on liver and lung peptide abundances at four gestational stages (E13, E14, E16, and P0). (**4 csv files**)
4. Proteomics data on the liver and lung at various gestational stages of mouse fetus. (**1 csv file**)

## Supporting information

supplementary data 2A brain rep1

supplementary data 2A liver rep1

## ACKNOWLEDGEMENTS

We would like to thank Lindsay K. Pino for her helpful suggestions. BEC was supported by a TL1 Award. Research reported in this publication was supported by the National Center for Advancing Translational Sciences of the National Institutes of Health under award number TL1TR001880. BAG gratefully acknowledges support from NIH grants NS111997 and HD106051. The content is solely the responsibility of the authors and does not necessarily represent the official views of the National Institutes of Health.

## AUTHOR CONTRIBUTIONS

JB and BEC conceived the paper and designed the experiments. BEC performed the fetal and neonatal injections. JB, JR and MM performed mass spectrometry and downstream data analysis. WHP and BAG guided experimental design and parameters. JB, BEC and ZL drafted the manuscript. ZL, WHP and BAG revised the manuscript and provided formative feedback.

## STAR METHODS

### KEY RESOURCES TABLE

| REAGENT or RESOURCE | SOURCE | IDENTIFIER |
| --- | --- | --- |
| **Deposited data** |  |  |
| Raw files and search results | This paper | MassIVE repository (identifier: MSV000087342) |
| **Software and algorithms** |  |  |
| R scripts | This paper | https://github.com/baezaj/in-utero-turnover |

### RESOURCE AVAILABILITY

#### Lead contact

Further information and requests for resources and reagents should be directed to and will be fulfilled by the lead contacts, William H. Peranteau, Benjamin A Garcia.

#### Materials availability

This study did not generate new unique reagents.

#### Data and code availability

- The data used in this study can be accessed at xxx.
- The source code can be accessed at https://github.com/baezaj/in-utero-turnover. DOIs are provided in the key resources table.
- Any additional information required to reanalyze the data reported in this paper is available from the lead contact upon request.

### METHODS DETAILS

#### Animals

C57BL/6J (B6, Jackson Cat. No. 000664) were purchased from Jackson Laboratories and a colony to produce time-dated pregnancies was maintained at our facility. Experimental protocols were approved by the Institutional Animal Care and Use Committee and followed guidelines outlined in the National Institutes of Health Guide for the Care and Use of Laboratory Animals.

#### *In utero* and neonatal pulse-injection of labeled amino acids

Fetuses of time-dated pregnant B6 mice were injected via the vitelline vein at gestational day (E) 13.5, 14.5, or 16.5 (as indicated in the Results sections) with labeled amino acids as previously described^20, 21^. Briefly, under isoflurane anesthesia (1% to 5%) and using a sterile technique, a midline laparotomy was performed and the uterine horns were exposed. Labeled amino acid solution (10 μL of L-Lysine Hydrochloride and L-Arginine Hydrochloride [Silantes GmBH, Product Number 211604102] reconstituted in sterilized phosphate buffer solution) was delivered to each fetus. All injections were performed with a programmable microinjector (IM-300 Microinjector; MicroData Instrument Inc, S. Plainfield, NJ) using a ∼100-μm beveled glass micropipette and a dissecting microscope to improve accuracy and verify the location of injection^20^. The microinjector provides a high level of administrative control when introducing the heavy amino acid solution. Successful injections were confirmed by visualizing the clearance of blood in the vitelline vein by the injectate. After injection, the uterine horn with the fetuses was returned to the peritoneal cavity and 2-layer abdominal closure was performed with 4-0 Vicryl suture (Ethicon). Dams recovered for 15 to 20 min under a heat lamp until they were ambulatory. Scheduled postoperative analgesia (oral meloxicam 2.5 mg/kg) was delivered daily for 48 hours when applicable. Isotopic amino acids circulate throughout the fetus and are transported to developing organs where they are used by the translational machinery for protein synthesis. At the designated time points after amino acid injection, fetal tissues including the brain, liver, lung, and heart were harvested. The tissues are then flash-frozen using liquid nitrogen and stored at −80 °C until further processing.

In a similar fashion to that described above, day of life 0 (P0) neonatal mice were injected with labeled amino acids via the facial vein using a ∼100-μm beveled glass micropipette, the programmable microinjector, and a dissecting microscope. For P0 injections, a total volume of 20 L was injected.

Standardized control over the pulse injection begins with the formation of the labeled amino acid solution. Isotopic amino acids are diluted in sterile buffered saline, created in a large batch, and aliquoted into single-use aliquots, which ensures that each fetus or neonate is delivered the same concentration of heavy-isotope label amino acids. This step reduces the injection-to-injection variability that may incur from unique sample creation with each experiment.

#### Sample processing

Tissue samples were resuspended in a denaturing buffer (6 M Guanidine-HCl, 100 mM TEAB pH = 8.5) and lysed by sonication. Protein estimation was performed using Bradford reagent using BSA for the standard curve. An equal amount of protein (10 - 30 μg) was resuspended to a final concentration of 1 μg/μL using 6 or 8 M Guanidine-HCl, 100 mM TEAB pH = 8.5, 15 mM DTT (Guanidine-HCl concentration ranged between 4.5 M and 6 M). Samples were denatured and reduced by incubating at 60 °C for 20 minutes while shaking at 1000 RPM. Cysteine alkylation was performed by adding 30 mM iodoacetamide and incubating for 20 minutes in the dark at ambient temperature while vortexing at 1000 RPM. Before proteolytic digestion, guanidine-HCl was diluted to 1 M using 25 mM TEAB pH = 8.5. Trypsin (Promega, Madison, WI) was added to a final ratio of 1:60 (trypsin: protein) and incubated at ambient temperature overnight. Following trypsin digestion, peptides were desalted using in-house desalting tips made from 3M™ Empore™ C18 extraction disks.

#### LCMS analysis

Peptides were analyzed with a Dionex UltiMate 3000 UHPLC system coupled to a high-resolution orbitrap mass spectrometer. A 25 cm fused silica capillary (75 µm ID, 360 µm OD) with an in-house pulled emitter tip was packed with ReproSil-Pur 3 µm diameter, 120 Å pore size C18 beads (Dr. Maisch GmbH). Solvent A was composed of LCMS-grade water and 0.1% formic acid. Solvent B was composed of 80% acetonitrile, 20% water (*v*/*v*), and 0.1% formic acid. Peptides were preloaded onto a µ-Precolumn cartridge (300 µm ID, 5 mm) packed with C18 PepMap 100, 5 µm diameter, 100 Å pore size C18 particles (Thermo Scientific). Peptides were loaded with a 5 µL/min flow rate using 2% acetonitrile, 98% water, and 0.05% trifluoroacetic acid and back-flushed with Solvent A onto the pre-equilibrated analytical column. Peptides were separated using a 182-min segmented gradient that consisted of an isocratic hold at 5% B for 2 minutes, a linear increase from 5% to 25% B for 144 minutes, and 25% to 40% B for 36 minutes. The column was washed with 90% B for 10 minutes and re-equilibrated with 5% B for 7 minutes. The flow rate for the analytical column was set to 400 nL/min.

The eluted peptides were ionized by electrospray ionization and analyzed on an Orbitrap Q Exactive HF-X mass spectrometer (Thermo Scientific) in a data-dependent mode. The precursor scan range of 350 to 1500 m/z was acquired in profile mode at 60K resolution (at 200 m/z) with automatic gain control of 1E6 ions for a maximum of 50 milliseconds. The top 30 precursors were selected for fragmentation with a 0.7 Th window, fragmented in the HCD cell using a stepped normalized collision energy (NCE) of 25.5, 27, and 30, analyzed in the orbitrap at 15K resolution, and acquired in centroid mode. Ion accumulation was set to a maximum of 1E5 or 40 milliseconds. The dynamic exclusion was set to 45 seconds.

#### Data analysis

The raw data was processed using Thermo Proteome Discoverer (version 2.3.0.523, Thermo Scientific). The raw spectra were searched using a combination of spectral library and database search strategies. Spectral library search was performed using MSPepSearch with the NIST spectral libraries^53^: Orbitrap HCD 20160923 v.1 (1127970 spectra) and ProteomeTools synthetic HCD 20170530 v.1 (696692 spectra)^54^ and validated using Percolator with a 1% FDR. The combination database search was performed using MS Amanda 2.0 and Sequest HT. Tryptic peptides with up to two missed cleavages were searched against a Mus musculus database (SwissProt database; TaxID - 10090; version 2017-10-25; 25,097 entries) as well as a common contaminants database with a precursor tolerance of 20 ppm and a fragment mass tolerance of 0.05 Da. Carbamidomethylation of cysteines (+57.0215 Da) was set to a fixed modification; oxidation of methionines (+15.9949 Da) and N-terminal protein acetylation (+42.0106 Da) were set to dynamic modifications. For the isotopic amino acid injected samples, Lys8 (+8.0142 Da) and Arg10 (+10.0083 Da) were also set to dynamic modifications. A reverse decoy database was used to determine a 1% false discovery rate (FDR) using Percolator^55, 56^ with peptide spectral match (PSM) validation based on *q*-value.

#### Mass isotopomer distribution analysis

We determined the fractional abundance of the isotopic amino acid pool using mass isotopomer distribution analysis^29, 30^. In this analysis, peptides that contain either two lysines or two arginines are used and we used different database search engines to identify a larger pool of peptides that fit these criteria (Proteome Discoverer 2.3, MaxQuant^57^, and pyQms^58^). For each database search engine, we used the default parameters for a 2-plex SILAC experiment and included Lys8 and Arg10 as variable modifications. Peptides containing two lysines or arginines will be present in one of three states: all light or unlabeled (LL), all isotopic or heavy (HH), or a mix of light and heavy (HL). The equation used to determine the fractional abundance of the isotopic amino acids is:

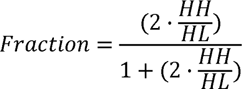

#### Determination of turnover rate

Heavy fractional abundance (H / (H + L)) for each peptide was calculated from the light and heavy abundance values. Turnover rates were determined by fitting the peptide fractional abundance as a function of time using a nonlinear least squares exponential function (NLS in R v4.0.2) using the following equation:

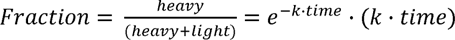

with *time* being the duration of time (in hours) between the pulse injection and when tissues were harvested. A linear model was used to estimate the starting parameters of the turnover model which was then allowed to run for 100 maximum iterations. The resulting turnover rate (*k*) for each peptide was used for downstream analysis or converted to half-life using the:

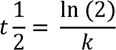

Turnover model coefficients were extracted using the broom package (v 0.7.4) in R and were filtered for *p* value < 0.05 and std. error < 0.025.

#### Statistical analysis

All statistical analyses were carried out in R Statistical software (v 4.0.2). *p*-values were adjusted for multiple hypothesis correction using the Benjamini-Hochberg method to report *q*-values.

## Supplementary Figures

**Figure S1:**
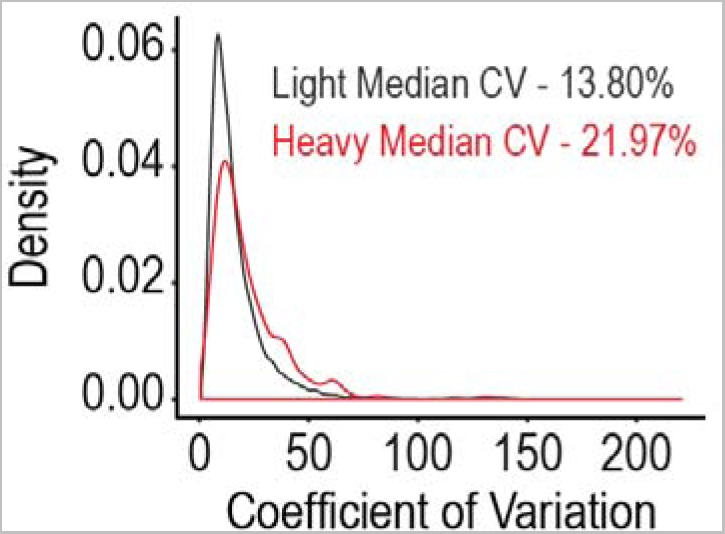
Peptide variability. Coefficient of variation (CV) for light and heavy peptides using six technical replicates. The coefficient of variation was calculated using normalized peptide abundances in the linear scale.

**Figure S2:**
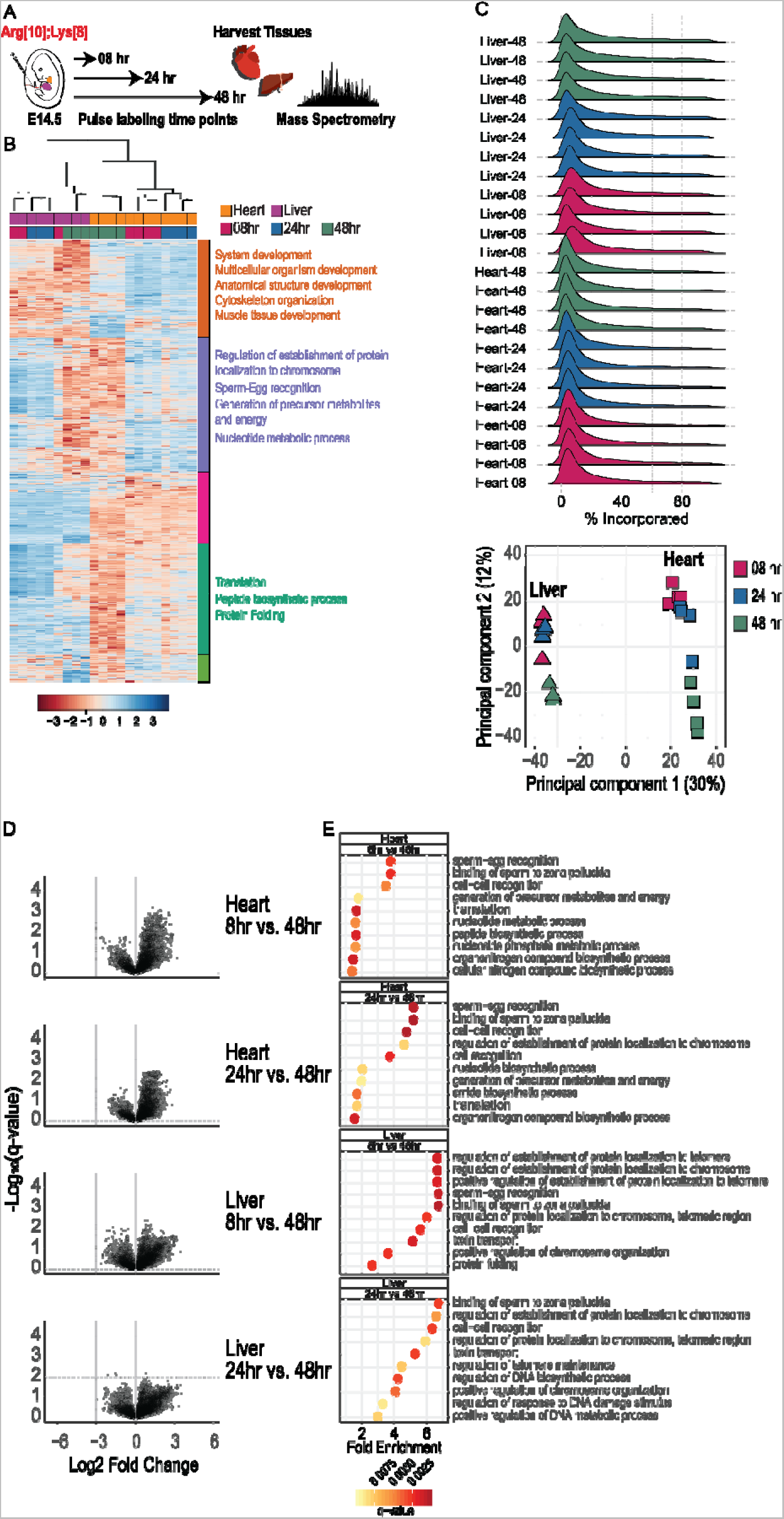
Labeling kinetics of labeled amino acids. A. Experimental design for assessing labeled amino acid incorporation. E14.5 fetal mice are pulse injected with labeled arginine (Arg10) and lysine (Lys8) followed by a labeling period of 8, 24, and 48 hours. Heart and liver tissues are harvested and analyzed by mass spectrometry. **B.** Heatmap of the heavy peptide abundances for all labeling periods in heart and lung tissues. **C.** Distribution of heavy amino acid incorporation for all labeling periods in the heart and liver. **D.** Volcano plot comparing the heavy peptide abundance between heart and liver. Statistical analysis was performed using a Student’s t-test and p.values were adjusted for multiple hypothesis testing. **E.** Gene ontology of the significant hits for each comparison.

**Figure S3:**
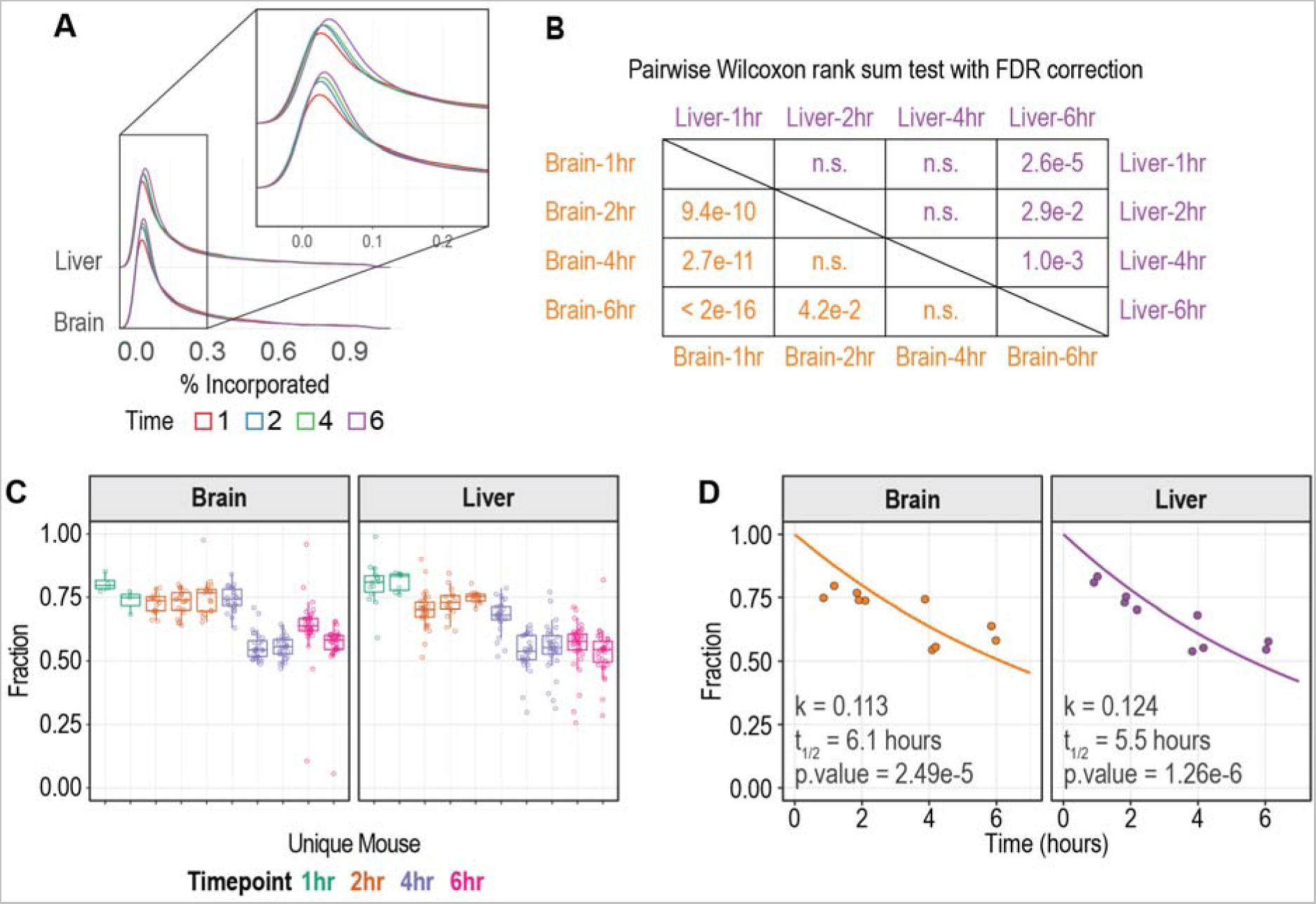
Validation of in utero turnover quantification. **A.** Distribution of heavy labeled amino acid incorporation into the fetal tissue proteome across the different time points. **B.** Pairwise comparison of the incorporation distribution from A. Wilcoxon rank sum test with FDR correction was used for the pairwise comparison. **C.** Relative isotope abundance (RIA) of the precursor amino acid pool quantified by mass isotopomer distribution analysis (MIDA). Peptides containing two lysines or two arginines for each sample were assessed to quantify the RIA as described previously^24, 30^. Individual points in the boxplots represent the calculated RIA value for unique peptides across samples and time points. **D.** The median RIA value from each sample in C. was used for a pulse-chase kinetic model.

**Figure S4:**
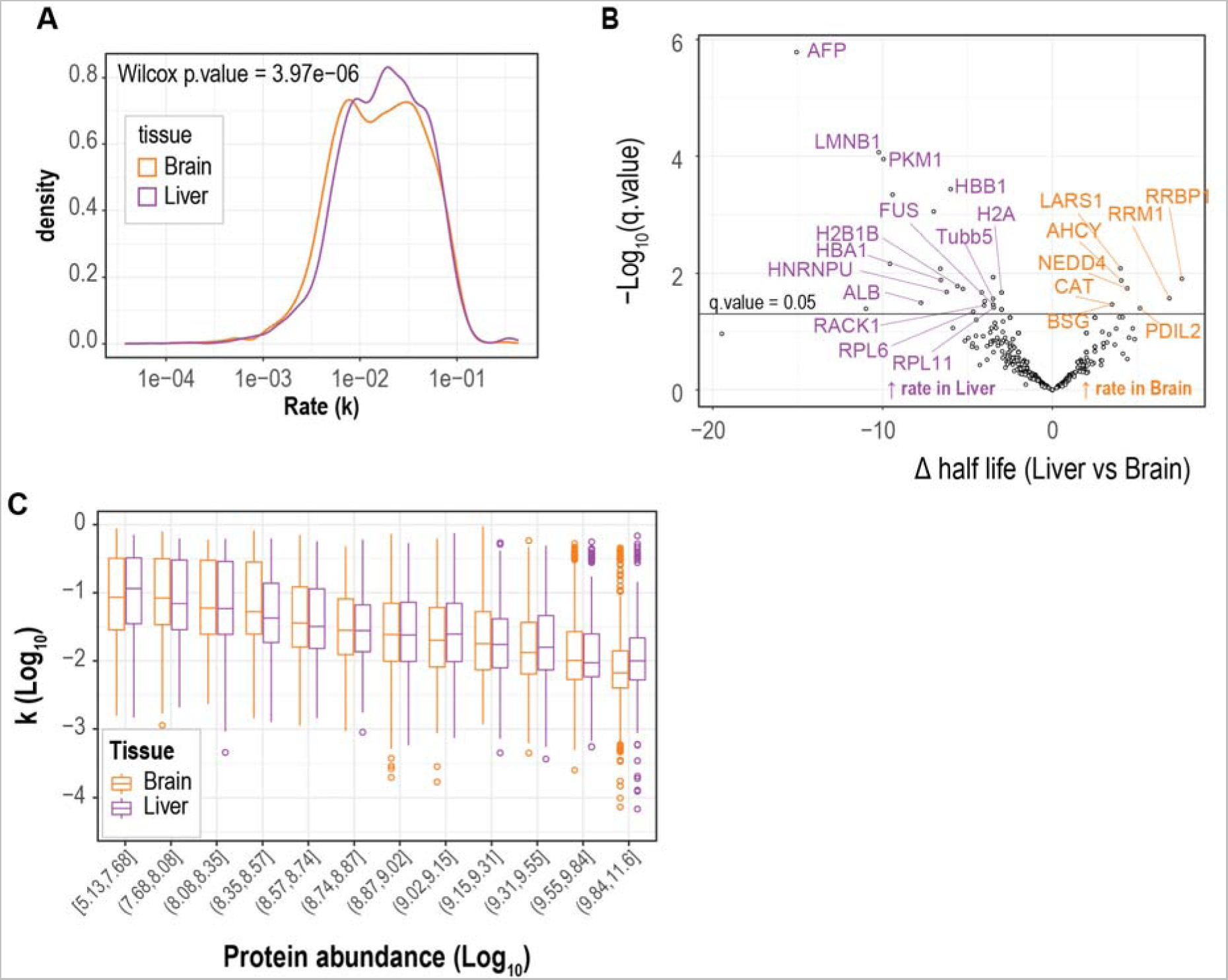
Liver and brain tissue turnover analysis. **A.** Distribution of quantified turnover rate in liver and brain tissues. A Wilcoxon rank sum test was used to compare the distributions. B. Volcano plot comparing half-lives of proteins between the liver and brain. The statistical test performed was a Wilcoxon rank sum test. **C.** Distribution of turnover rates as a function of protein abundance. Protein abundance is binned using groups with an approximately equal number of observations.

**Figure S5:**
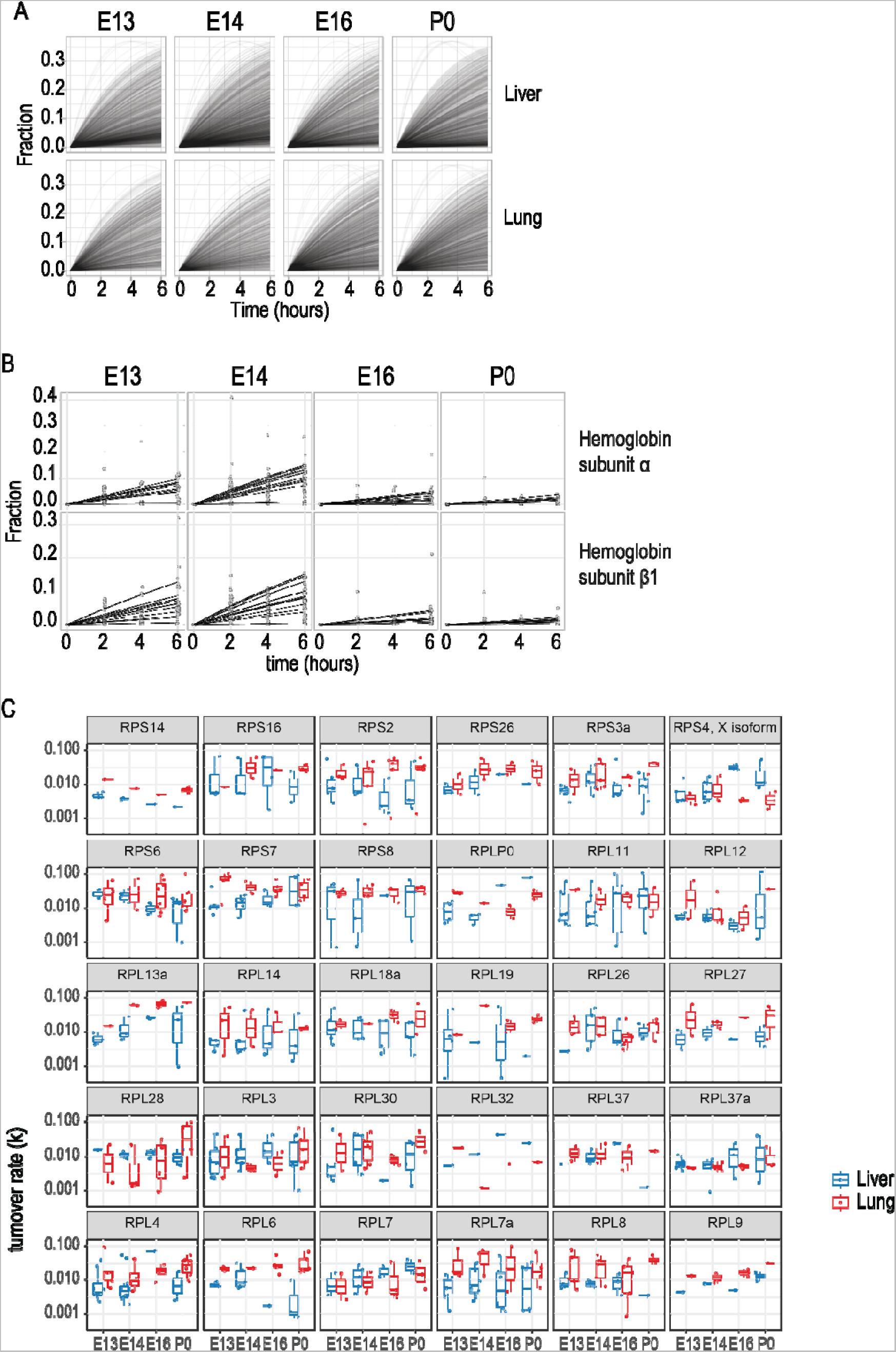
Quantifying turnover rates across mouse fetal development. **A.** Turnover rate profiles of all peptides for each gestational age and tissue. **B.** Turnover rate profiles showing all the data points and best-fit line for hemoglobin α and β1 in liver tissue. **C.** Summarized turnover rates quantified for ribosomal proteins shared between the liver and lung.

**Figure S6:**
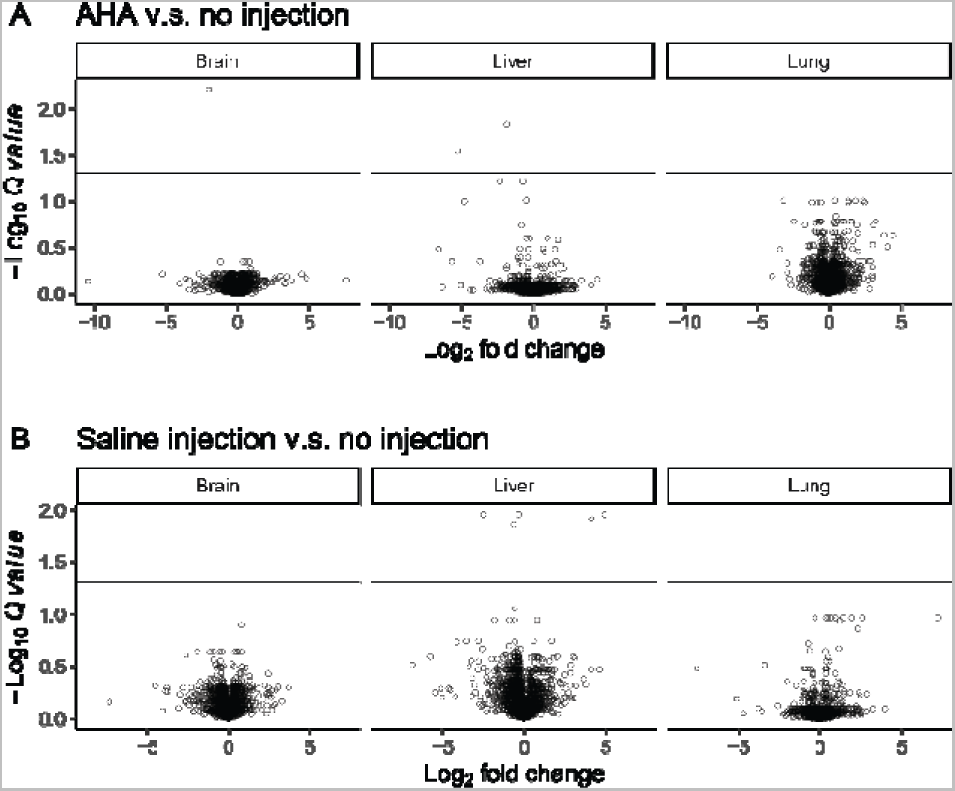
Volcano plots of three organs after AHA and saline injections. **A.** Volcano plot comparing the the proteomes after AHA injection and control (no injection). **B**. Volcano plot comparing the the proteomes after saline injection and control (no injection). Statistical analysis was performed using a Student’s *t*-test and *p* values were adjusted for multiple hypothesis testing.

